# Untargeted metabolomics reveals farming practice- and cultivar-driven modulations of pea (Pisum sativum L.) seed metabolome

**DOI:** 10.1101/2025.06.23.661072

**Authors:** Sarvar A. Kakhkhorov, Søren Balling Engelsen, Kristian Holst Laursen, Bekzod Khakimov

## Abstract

This work presents untargeted LC-MS-based metabolome of ten cultivars of peas grown at three field sites following different agricultural practices. More than 1,200 metabolite features were detected in methanolic extracts of pea seed flours. Of these, nearly 300 features were identified using mass spectral libraries and advanced computational tools. Approximately 40 metabolites were found to be associated with location effect, independent of cultivar type. Organically grown pea samples showed lower level of the main pea triterpene glycoside soyasaponin I and higher levels of nitrogen-abundant amino acids indicating increased nitrogen availability in soil. More than 100 metabolites were associated with the location-independent cultivar effect. Two cultivars resistant to downy mildew and pea wilt, Akooma and Greenway, showed the most distinct metabolome with greater levels of polyunsaturated fatty acids and lipid oxidation products known to give ‘beany’ off-flavors. The most commonly cultivated pea variety, Ingrid, had significantly lower levels or was completely devoid of hydroxycinnamic acid amides such as caffeoyl, feruloyl, and coumaroyl aspartates that were present in all other cultivars. Three chloroauxin metabolites, reported here for the first time, were identified through molecular networking within GNPS platform and propagation of annotation from a computationally predicted indole-3-acetic acid catabolite. Overall, the results indicate biochemical adaptation of pea plants to location or agricultural practice as reflected in their seed metabolome.

## Introduction

The awareness of sustainable food production is increasing due to the growing global population and climate change, along with a rising recognition of the importance of a balanced diet for human health (Khakimov et al., 2025). Considering these factors, more attention is being directed toward expanding the research on various crops to diversify dietary options while promoting both health and environmental resilience (Dwivedi et al., 2017). In this regard, pulses are nutrient-dense staple crops serving as valuable sources of protein, dietary fiber and various carbohydrates, vitamins, mono- and polyunsaturated fatty acids, and essential micronutrients in the human diet. Besides the major staple crops such as wheat, rice, maize, and soybean, the pea seeds (*Pisum sativum L.*) provide notable agricultural and nutritional benefits. Although the starch content of dried pea seeds ranges between 40-60%, their glycemic index is relatively lower than that of most carbohydrate-rich crops, making them a calorific yet nutritionally healthier option (Mudryj et al., 2014). As members of the nitrogen-fixing *Fabaceae* family, pea seeds are rich in protein and comprises an overall average of 23.4% of their dry mass (Mudryj et al., 2014). Glutamine, aspartic acid, arginine, and lysine were found to be among the most abundant amino acids in peas, whereas methionine, tryptophan, and cysteine were present at lower levels (Tömösközi et al., 2001). The same study showed that pea seeds outperformed lupin and soybean in the levels of essential amino acids such as arginine, valine, and methionine. The lipid content of peas ranges from 0.9 to 5.0 % and the amount of polyunsaturated fatty acids (PUFA) is higher than that of saturated fatty acids (Khodapanahi et al., 2012).

As sessile organisms, plants have developed a remarkable genetic capacity to produce a diverse array of specialized metabolites that enable them to adapt to environmental challenges (Weng et al., 2021). While these small molecules serve critical roles in plant defense and adaptation, they are also of significant interest in the context of plant-based foods due to their effects on nutritional value, sensory properties, and potential health benefits. Pea seeds, for instance, are particularly rich in phytochemicals such as phenolic compounds, carotenoids, and saponins (Dahl et al., 2012). In addition, several glycosides of quercetin, apigenin, luteolin, kaempferol, myricetin, and isorhamnetin have been identified in pea seeds and are known to contribute to their antioxidant and anti-inflammatory properties (Lim, 2012; Wu et al., 2023). Like many legumes, pea seeds contain various triterpene glycosides, such as group B saponins (soyasaponin Ba, Bb, Bx, Bc) and DDMP (2,3-dihydro-2,5-dihydroxy-6-methyl-4H-pyran-4-one) saponins (soyasaponin αg, βg, γg, αa, βa), which are considered to account for their bitterness and astringency (Heng et al., 2006). Despite the concentrations of these secondary metabolites are relatively small in pea seeds, they play a crucial role in the digestibility of dominating macromolecules like pea protein and fiber and contribute to an overall nutritional value of pea (Hao et al., 2022). It is therefore important to map not only macromolecules but also small molecular metabolites in pea shaping its chemical space to better understand their nutritional value, sensory characteristics, and potential impact on human health. Moreover, the variation in cultivar type and growing conditions, including geographical location and agricultural practices, influences biosynthetic pathways in pea, resulting in differences in their metabolome. Studies have shown variation in the profiles of amino acids, lipid-like molecules, polyketides, and phenylpropanoids in eight pea cultivars grown in different regions of Spain using the combination of targeted and untargeted LC-MS metabolomics (Reveglia et al., 2025). Witten et al. reported that crude protein content was negatively correlated with the essential amino acids contents in various organically grown cereals and legumes including peas (Witten et al., 2020). The same study also demonstrated that harvest site and time of the year had greater impact on nutrient composition (protein and amino acid content) than the crop variety. Lastly, in a study of two pea cultivars grown under normal condition and biotic stress represented as contamination with soil-borne pathogens, cultivar effect was shown to influence volatile compounds and lipoxygenase activity (Oliete et al., 2022). This enzyme leads to lipid peroxidation of PUFAs as a response to pathogen attack that eventually contributes to the accumulation of off-flavor substances in pea seeds.

Several analytical methods have been used to examine the chemical profile of crops, both qualitatively and quantitatively (Bertram et al., 2010; Bljahhina et al., 2023; Khakimov et al., 2017; Khakimov et al., 2016). In this regard, untargeted metabolomics facilitates an unbiased analysis of all detectable metabolites with molecular masses ≤1,500 Da in biological systems. Combined with chemometrics, it uncovers distinct molecular patterns associated with one or more factors included in a study design (Khakimov et al., 2017). Ultra-high performance liquid chromatography (UPLC) coupled with high-resolution mass spectrometry (HRMS) has become the method of choice in metabolomics due to its broad metabolite coverage, high sensitivity, and selectivity. When combined with emerging computational mass spectrometry approaches (Dührkop et al., 2019; Wang et al., 2016) liquid chromatography–tandem mass spectrometry (LC-MS/MS) enables comprehensive profiling of small molecules in biological systems, making it a powerful tool in metabolomics and foodomics research (Christ et al., 2018; Gauglitz et al., 2020; Gauglitz et al., 2022; Khakimov et al., 2015).

In this study, untargeted LC-MS/MS-based metabolomics was employed to investigate the metabolite variation in pea seed flours from ten cultivars grown across three geographic locations in Denmark, including two farming sites managed conventionally and one site managed under organic agriculture practices. Mass spectral libraries along with advanced, mass spectral computational tools for molecular networking in Global Natural Products Social Molecular Networking (GNPS) platform paired with molecular structure identification in SIRIUS were employed to annotate metabolites in LC-MS/MS data. Using the propagation of annotation in molecular networking within GNPS platform, the structure of three chloroauxins is putatively annotated for the first time. Application of chemometrics on metabolomics data revealed the distinct effect of growth location and pea cultivar, and their interaction on pea seed metabolome.

## Results

### Pea metabolome

The final feature table included 1230 variables after Metaboscape-based processing of raw LC-MS/MS data files obtained on 90 pea samples (3 locations x 10 cultivar x 3 replicates). Using mass spectral libraries implemented in Metaboscape, 121 features were identified at Level 2 in accordance with the Metabolomics Standards Initiative (Sumner et al., 2007) (Supplementary Table 1). Similarly, feature-based molecular networking (FBMN) using publicly available spectral libraries within the GNPS platform allowed annotation of 173 features (Nothias et al., 2020) (Supplementary Table 1). 118 GNPS annotations were derived from the nearest-neighbor suspect spectral library, which was generated through the propagation of annotations of reference compounds in repository-scale molecular networking (Bittremieux et al., 2023). Additionally, 178 metabolites were annotated using molecular formula and structure identification tools CSI(Compound Structure Identification):FingerID (Dührkop et al., 2015; Hoffmann et al., 2022) in SIRIUS platform. In total, the number of unique annotations from spectral libraries in Metaboscape and GNPS was 30 and 70, respectively, while 18 features were dereplicated using these two annotation sources (Figure 1). The number of shared annotations between SIRIUS and spectral libraries in Metaboscape and GNPS were 33 and 45, respectively, whereas 60 features were annotated using SIRIUS only. Lastly, 40 LC-MS/MS features could be annotated by all three sources and 948 features remained non-annotated. Further, compound class prediction using CANOPUS in SIRIUS platform was performed to assign molecular class to 424 metabolites representing 42 molecular classes (Dührkop et al., 2021). Major molecular classes included 126 fatty acid derivatives (amides, acyls, esters, glycosides, octadecanoids), 74 amino acids, small peptides, and oligopeptides, 71 glycero(phospho)lipids, 44 alkaloids (pseudo-, tryptophan, histidine, nicotinic acid, tyrosine, anthranilic acid), 23 terpenoids (sesqui-, tri-, di-, mono-), 17 nucleosides, 11 phenylpropanoids (C6-C3), 9 flavonoids and phenolic acids, and 9 saccharides (Supplementary Table 1). The downstream data analysis was performed separately (1) for 121 metabolites annotated in Level 2 using Metaboscape, (2) for 178 metabolites annotated in Level 3 using SIRIUS, and (3) for all 1230 variables including 948 unknown features.

**Figure 1.**
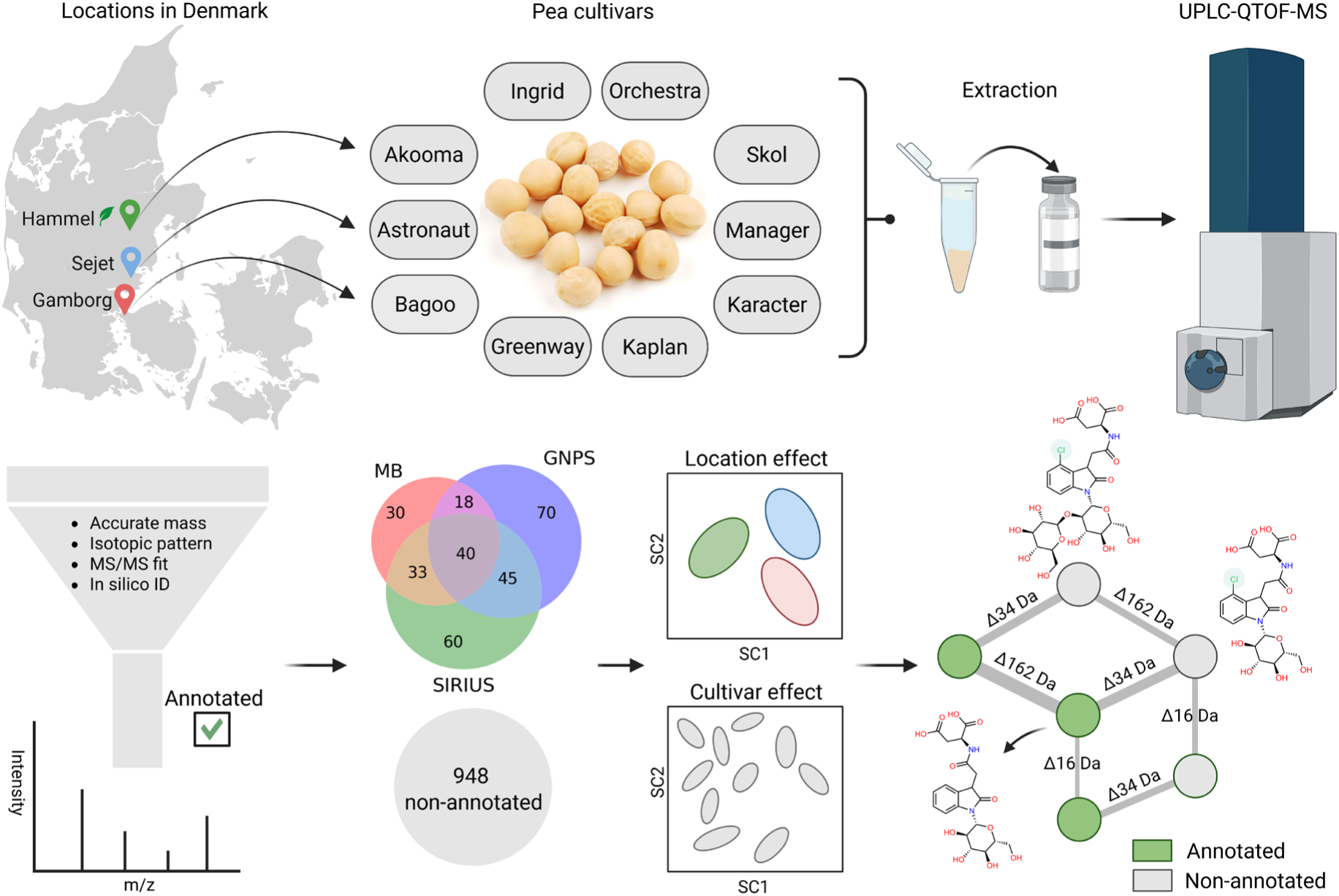
UPLC-QTOF-MS-based untargeted metabolomics study of ten pea cultivars grown in three different locations in Denmark with organic agriculture practice used in Hammel. In total, 90 samples (10 cultivars × 3 locations × 3 replicates) along with pooled quality control samples were extracted in 80% methanol with addition of internal standard to acquire untargeted MS data. Raw data were processed and annotated using spectral libraries and computational approaches with pre-defined parameters, which was then analysed using ANOVA-simultaneous component analysis (ASCA) to reveal the effect of location, cultivar, and their interaction on pea metabolome. The putative annotation of significant non-annotated features was additionally revealed using computational approaches GNPS and SIRIUS.

### ASCA-based decomposition of variation in data according to the study design

The full-factorial design of the study, including two factors: 1) location with three levels and 2) cultivar with ten levels, allowed the use of ASCA with permutation testing (n=10,000) to investigate the variation present in the data that derive from the two factors, and their interaction terms on pea metabolome (Figure 2). Both factors, location and cultivar, and their interaction effect were shown to be statistically significant when considering the 121 metabolites identified on the Level 2 annotation confidence, with highest variation attributed due to the cultivar effect (28.6%, p = 0.001), followed by the two-factor interaction effect explaining 17.2% of variation (p = 0.02). The location effect explained the lowest variation of 7.8% (p = 0.001) in the Level 2 metabolome (121 metabolites), despite organic agriculture being practiced in only one of the sites - Hammel. This trend was consistent when the Level 3 metabolome (178 metabolites) or the entire dataset with all 1230 features were analyzed using ASCA (Supplementary Figure 1-2).

**Figure 2.**
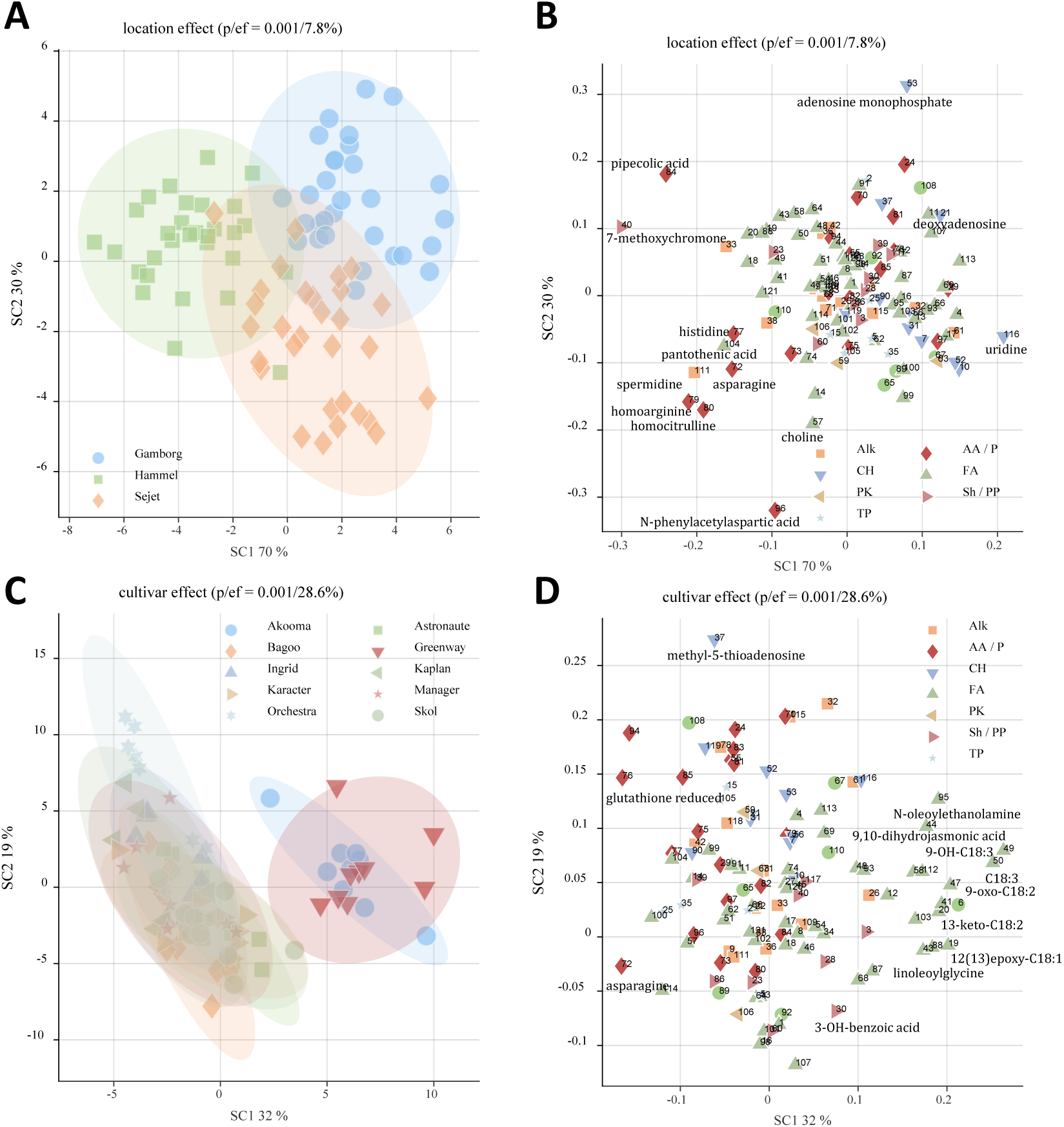
ANOVA-simultaneous component analysis (ASCA)-based location and cultivar effect. (A) Scores plot demonstrates partial separation of pea samples according to the three locations (variance explained = 7.8%, p-value = 0.001) (n = 3 × 30) with more prominent differentiation of pea grown at organic farming side Hammel. Loadings plots show the distribution of metabolites, coloured according to biosynthetic pathways they derived from, explaining location (B) and cultivar (D) effect. (C) Scores plot highlights cultivar effect (variance explained = 28.6%, p-value = 0.001) for all pea samples from three locations (n = 10 × 9) with Akooma and Greenway being most differentiated from other cultivars due to polyunsaturated fatty acids profile.

### Effect of location

As illustrated in the ASCA scores plot (Figure 2A), the main variation attributed to the location effect was driven by metabolome differences between the samples from Hammel and those from the other two sites, Gamborg and Sejet. Simultaneous component 1 (SC1) primarily differentiate the Hammel samples from those grown in Gamborg and Sejet, while SC2 differentiate the Gamborg and Sejet samples. The major overlap observed along SC1 and partial overlap observed along SC2 suggests a subtle dissimilarity in their metabolite profiles. Given that peas in Hammel were cultivated organically, whereas those in Gamborg and Sejet were conventional grown, the clear separation along SC1 largely reflect metabolome differences arising from agricultural practices. In this regard, ASCA loadings plot demonstrated the distinctive patterns of metabolites distinguishing the samples from the three locations (Figure 2B). Specifically, the metabolite profile distinguishing Hammel samples included increased levels of several amino acids—pipecolic acid, histidine, asparagine, homoarginine, and homocitrulline—as well as lipid-like compounds such as 13-keto-9,11-octadecadienoic acid, 1-pentadecanoyl-sn-glycero-3-phosphocholine, and pantothenic acid. In contrast, Gamborg samples were characterized by higher levels of adenosine 5′-monophosphate (AMP), 2-deoxyadenosine, and glycerophosphocholine, while Sejet samples were richer in N-phenylacetyl aspartic acid, choline, and LysoPC(14:0/0:0). Additionally, indole-3-lactic acid (ILA), uridine, oxidized glutathione, 3-aminobenzoic acid, and 1,6-anhydro-β-glucose were more abundant in samples from Gamborg and Sejet compared to Hammel.

The ASCA analysis performed on the Level 3 identified metabolites, including 178 metabolites annotated via SIRIUS (Supplementary Figure 1A), revealed a similar trend in the location effect, which accounted for 7.3% of the total variation present in the data (p-value=0.001). Among the computationally annotated metabolites differentiating the Hammel samples from those of the other two locations were pisatin, a common pea phytoalexin, argininosuccinic acid, and several fatty acids.

Sejet-grown peas exhibited relatively higher levels of C5:0-arginine (C5:0-Arg) and N-salicyloyl aspartic acid, while the Gamborg samples were characterized by elevated levels of triterpenoid saponins, including pisumsaponin I and soyasaponin I, as well as indole-3-acetyl aspartic acid (IA-Asp), a phytohormone belonging to the auxin class (Supplementary Figure 1B).

When all 1230 features were subjected to ASCA, the location effect again consistently explained 6.4% of the total variation (p-value = 0.001). The ASCA scores plot depicted the partial separation of Hammel samples from those grown in Sejet and Gamborg (Supplementary Figure 2A). Notably, samples from all three locations were split into two subpopulations, differentiating along SC1 (73%) and SC2 (27%). The two sub-populations within each location represented distinct cultivar types, and this distinction was consistent across the three locations. Although the interpretation of the ASCA loadings plot for 1230 features is inherently challenging, metabolites such as AMP, pisumsaponin I, benzoyl-Asp, 4-hydroxyproline, C5:0-Arg, and N-phenylacetyl-Asp contributed notably to the differentiation of location types (Supplementary Figure 2B).

To assess how individual metabolites vary across the three locations, independent of cultivar types, a one-way ANOVA was performed with Benjamini-Hochberg correction for multiple testing (FDR = 5%). Of the 1230 features analyzed, 205 showed significant differences in levels across locations. Among these, 33 out of 121 Level-2 identified metabolites differed significantly among pea seeds from the three locations. Pipecolic acid exhibited the largest effect size (η^2^) of 47.8% (p = 3.89e-11), being more abundant in Hammel samples compared to the other two conventional farming sites, Gamborg and Sejet (Figure 3B). Homoarginine (η^2^ = 36.6%, p = 1.28e-07), homocitrulline (η^2^ = 32.4%, p = 1.28e-07), and spermidine (η^2^ = 31.1%, p = 2.02e-06) showed a similar trend, being most abundant in Hammel, followed by Sejet and Gamborg samples. Conversely, N-phenylacetyl aspartic acid (η^2^ = 35%, p = 2.71e-07) was most abundant in Sejet samples, followed by Hammel and Gamborg, while adenosine 5-monophosphate (η^2^ = 32%, p = 1.31e-06) demonstrated the reverse trend. Uridine was notably less abundant in most Hammel samples (η^2^ = 29.8%, p = 3.94e-06), whereas Gamborg and Sejet samples showed comparably higher levels. The remaining significant metabolites showed effect sizes ranging from 10 to 19% (data not shown). Among them, two other nucleosides, adenosine and 2-deoxyadenosine, showed accumulation trend across locations similar to uridine. Further, amino acids (histidine and asparagine), vitamin B5 (pantothenic acid), lipid oxidation products (13-keto-octadecadienoic acid, epoxy-octadecenoic acid) were more abundant in Hammel samples compared to those from Gamborg and Sejet.

**Figure 3.**
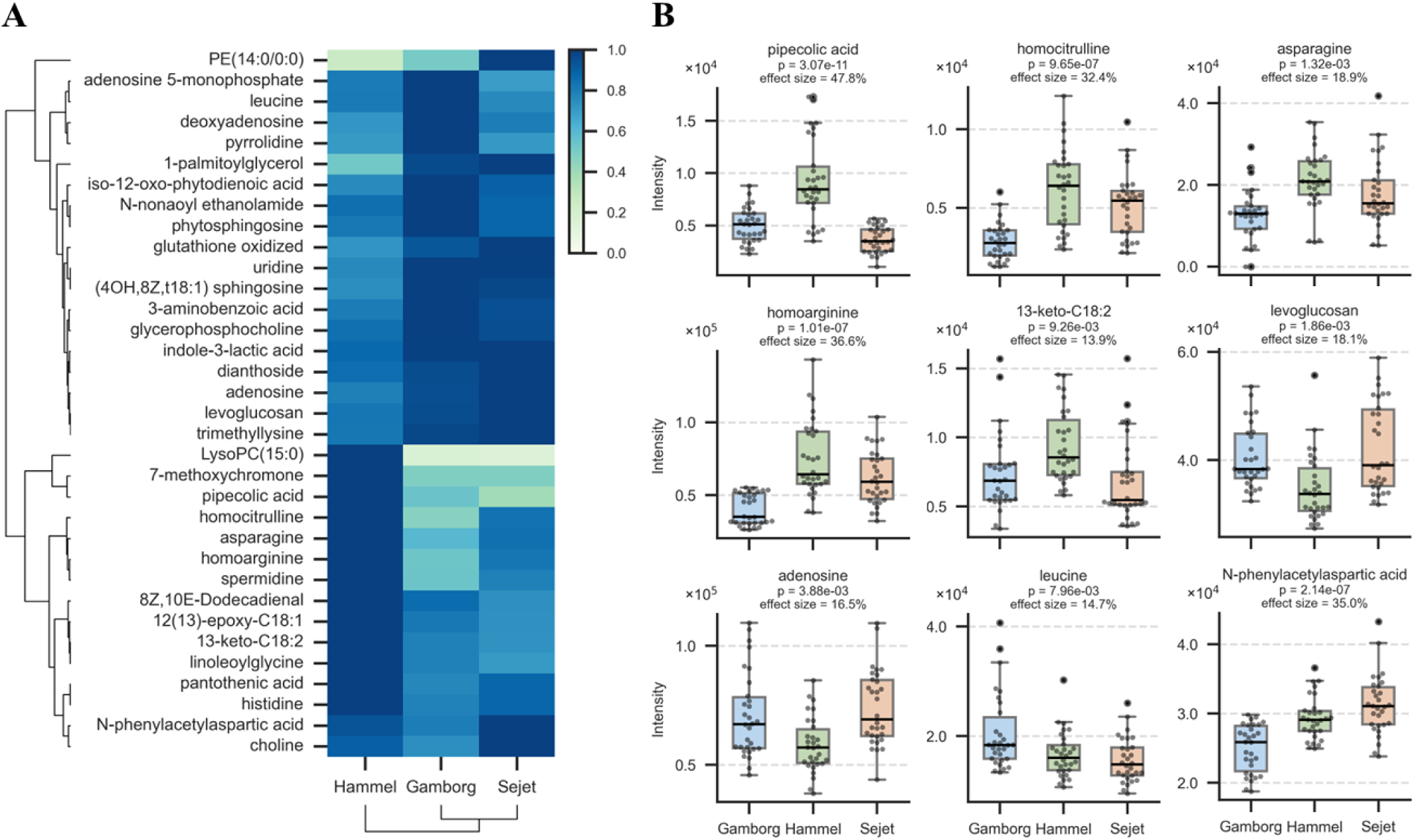
Cultivar-independent variation of metabolites levels annotated by spectral libraries across three locations. (A) Hierarchically clustered heatmap of 34 metabolites identified by one-way ANOVA (FDR-adjusted p-values < 0.05 using Benjamini-Hochberg’s correction for multiple testing) as significantly different between the three locations. (B) Box plots of selected metabolites with distinct relative abundance across locations identified as significant by one-way ANOVA.

The 33 metabolites that significantly differ between the three locations were visualized using a hierarchically clustered heatmap (Figure 3A). Hierarchical clustering revealed two main clusters: one comprising metabolites predominantly abundant in Hammel samples and another showing metabolites abundant in Gamborg and Sejet samples. Notably, amino acids with elevated abundance in Hammel such as homocitrulline, asparagine, homoarginine, and polyamine spermidine exhibited similar intensity patterns and were combined into one group, while histidine, pantothenic acid, and lipid-like compounds 13-keto-octadecadienoic acid, epoxy-octadecenoic acid, and linoleoylglycine formed another group that were also dominated in pea from Hammel.

Among the 178 metabolites annotated using SIRIUS, a one-way ANOVA identified 41 metabolites that are significantly influenced by location (FDR-adjusted p-values < 0.05) (Supplementary Figure 3B). Of these, argininosuccinic acid, a precursor to arginine in plants, exhibited the largest effect size of 52.1% (p = 2.31e-12), with the highest abundance in Hammel samples, followed by Sejet and Gamborg. Pisatin (η^2^ = 41.5%, p = 2.62e-09) showed a similar pattern. Additionally, the feature annotated as C5:0-Arg (using both the recent acyl amides library in GNPS and SIRIUS) had an effect size of 42.8% (p = 1.22e-09) and was slightly more abundant in Sejet samples compared to Hammel, with Gamborg samples showing the lowest levels. Among other significant metabolites annotated exclusively by SIRIUS, N-salicyloyl aspartic acid (η^2^ = 19.3%, p = 1.06e-03) was more abundant in Sejet samples, while indole-3-acetyl-N-aspartic acid (IA-Asp) (η^2^ = 16.8%, p = 3.00e-03) with slightly lower in organically grown Hammel samples. Pisumsaponin I (η^2^ = 10.7%, p = 3.29e-02) showed marginally higher levels in Gamborg and Hammel samples. Interestingly, soyasaponin I (η^2^ = 11.7%, p = 2.36e-02) showed its lowest abundance in Hammel samples, contrary to other secondary metabolites known for plant defense mechanisms, such as pisatin and pisumsaponin I.

When visualizing all location-influenced SIRIUS annotated metabolites in a heatmap, two distinct clusters emerged. The first cluster included 21 metabolites predominantly abundant in Gamborg and Sejet samples compared to samples from the organic farming site at Hammel. The second cluster included 17 metabolites with greater accumulation in Hammel samples (Supplementary Figure 3A). The pea phytoalexin pisatin grouped together with argininosuccinic acid, homocitrulline, asparagine, homoarginine, and spermidine in the second cluster. These compounds then clustered together with C5:0-Arg, which was most abundant in Sejet samples and slightly less abundant in Hammel samples. Additionally, N-salicyloyl aspartic acid, N-phenylacetyl aspartic acid, and choline formed a group showing similar abundance patterns, being more concentrated in Sejet samples than in Hammel and Gamborg samples. Pisumsaponin I group with pentadecanamide and cinnamic acid, while soyasaponin I was associated with indole-3-lactic acid. Finally, indole-3-acetyl-aspartic acid, a metabolite associate with the indole-3-acetic acid inactivation pathway, group with uridine and sphingosine.

### Effect of cultivar

The cultivar effect explained 28.6% (p = 0.001) of the variation in pea seed metabolome data, consisting of 121 Level 2 identified metabolites, as determined by ASCA. The ASCA scores plot indicated a clear separation of Akooma and Greenway cultivars from the remaining eight cultivars along SC1 (Figure 2C). The corresponding loadings plot suggested that these two cultivars contained relatively higher levels of fatty acids and their derivatives (Figure 2D). Specific discriminative markers for Akooma and Greenway cultivars included oxidation products of polyunsaturated fatty acids (C18:n), such as 9-oxo-octadecadienoic acid, 9-hydroxy-octadecatrienoic acid, octadecynoic (stearolic) acid, and 8,10-octadecadienal, that were found in relatively higher concentrations in these two cultivars. The remaining eight cultivars were partially separated along SC2, though the location effect seemed to remain since replicates of cultivars were further separated from each other depending on the farming site. Hammel-grown Orchestra, Kaplan, and Ingrid, the latter being a popular variety grown in Denmark, were most different from the other cultivars, having higher positive scores on SC2, and characterized by having greater levels of 5-methylthioadenosine, histidinol, N-benzoyl aspartic acid, and reduced glutathione (GSH).

The ASCA conducted on 178 metabolites identified using SIRIUS revealed a significant cultivar-influenced metabolic variation, accounting for 30% of the total variation (p = 0.001). The ASCA scores plot displayed a clear separation of Akoma, Greenway, and Ingrid cultivars, and partial separation of Orchestra, along SC1 (Supplementary Figure 1C). These four cultivars were further differentiated from the remaining six cultivars along SC2. Notably, cinnamic acid derivatives such as caffeoyl-, coumaroyl-, and feruloyl aspartates contributed for differentiation along SC2. Carboxylic acid conjugates of aspartate and glutamate (Asp-C4:0 and Glu-C5:0, respectively), indole-3-acetyl-N-aspartic acid (IA-Asp), 9-oxo-octadecadienoic acid, 9-oxo-octadecatrienoic acid, and soyasaponin I was the main metabolites responsible for the separation along SC1.

The ASCA performed on the entire dataset, comprising 1230 features, demonstrated an even stronger separation among cultivars, which explained 19.2% of total variation (p = 0.001) (Supplementary Figure 2C). Notably, Orchestra, Kaplan, Ingrid, Manager, Bagoo, Skol, and Karacter cultivars formed two distinct populations along SC2, indicating an impact of location which differentiates peas from the same cultivar. The ASCA loadings plot illustrates that Glu-C5:0, Asp-C4:0, benzoyl-Asp, and 5-methylthioadenosine are the key drivers for this clustering pattern (Supplementary Figure 2D). Interestingly, Akooma and Greenway cultivars again formed a distinct cluster separated from the other cultivars. Similarly, lipid oxidation products and other lipid-like compounds emerged as primary drivers for differentiating Akooma and Greenway cultivars from the rest. Among the prominent non-annotated features, metabolites belonging to polyketides, alkaloids, and terpenoid pathways differentiate Akooma and Greenway samples.

One-way ANOVA revealed that 74 out of 121 Level 2 identified metabolites exhibited significant difference among pea cultivars, independent of the location effect. The effect sizes ranged from 20.3% to 92.5%. Chromone showed the highest effect (92.5%, p = 4.9e-39), being most abundant in Orchestra samples, followed by Karacter, and absent in Ingrid (Figure 4A). Ingrid samples also displayed negligible levels of protocatechuic acid (η^2^ = 80.1%, p = 9.80e-23), a metabolite previously correlated with antioxidant capacity in peas (Chen et al., 2023).

**Figure 4.**
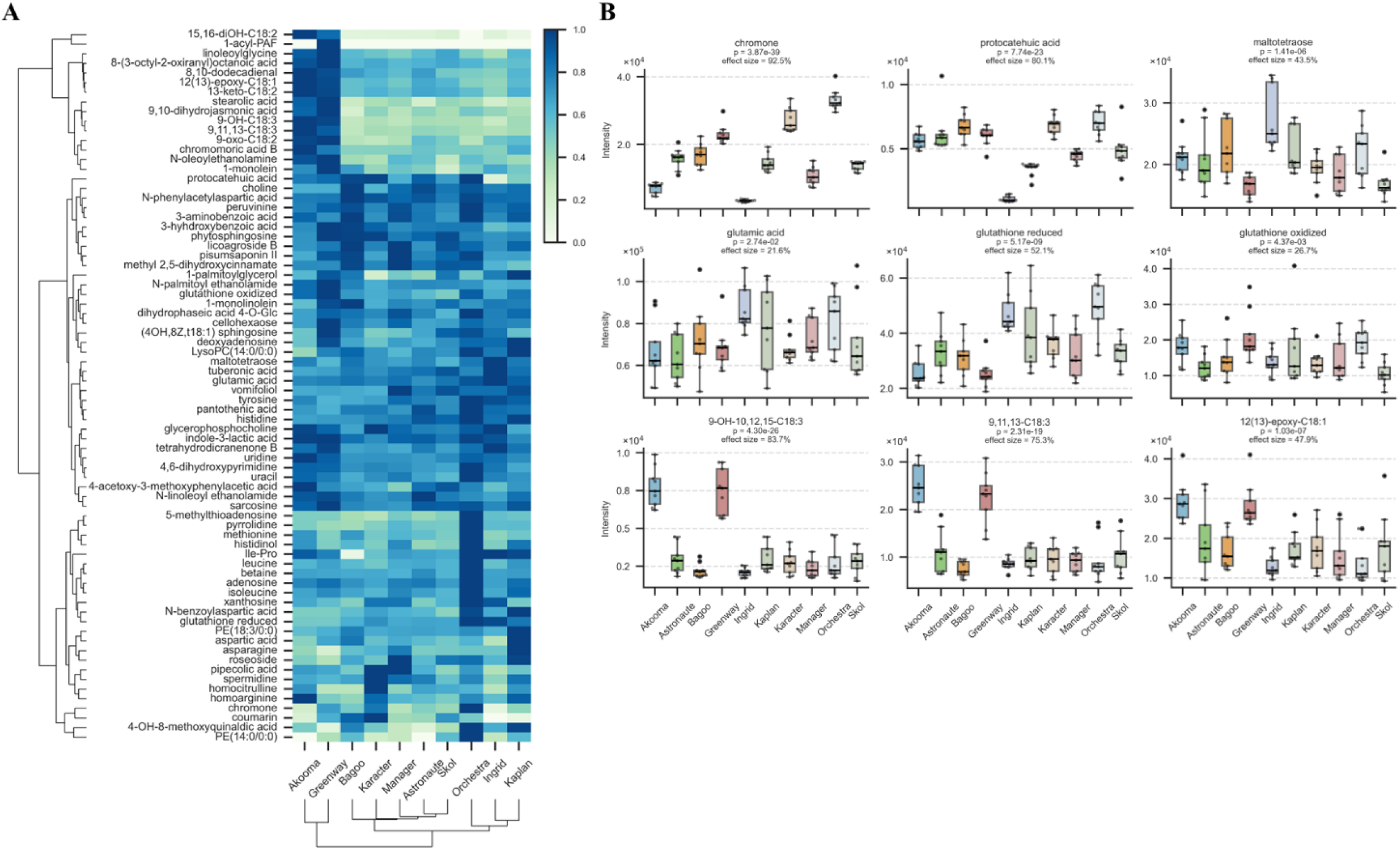
Location-independent variation of metabolites levels annotated by spectral libraries across ten cultivars. (A) Hierarchically clustered heatmap of 74 metabolites identified by one-way ANOVA (FDR-adjusted p-values < 0.05 using Benjamini-Hochberg’s correction for multiple testing) as significantly different between the cultivars. (B) Box plots of selected metabolites with distinct relative abundance across cultivars identified as significant by one-way ANOVA.

Akooma and Greenway cultivars contained the highest levels of polyunsaturated fatty acid (PUFA)-derived oxylipins, including 9-hydroxy-octadecatrienoic acid (η^2^ = 83.7%, p = 5.44e −26), 9-oxo-octadecadienoic acid (η = 76%, p = 1.10e-19), and octadecatrienoic acid (η^2^ = 75.3%, p = 2.34e-19). Regarding amino acids, Kaplan and Bagoo cultivars showed the highest abundance of aspartic acid (η = 66.2%, p = 2.76e-14), while Ingrid exhibited the lowest. Kaplan samples were also notably richer in asparagine (η^2^ = 59.5%, p = 2.35e-11), with Greenway having the least abundance. Homocitrulline levels were lowest in Bagoo, Greenway, and Ingrid cultivars (η = 52.6%, p = 5.08e-09). Although homocitrulline is associated with urea cycle disorders in humans, its biological role in peas remains unexplored (Turunen et al., 2014). Reduced glutathione was abundant in Ingrid, Kaplan, and Orchestra samples (η^2^ = 52.1%, p = 6.55e-09). Leucine (η^2^ = 36.4%, p = 6.62e-05) and 5-methylthioadenosine, a metabolite involved in the plant methionine salvage pathway (η^2^ = 58.6%, p = 5.08e-11), were highest in Orchestra samples, with other cultivars showing comparable levels. Methionine (η^2^ = 53.3%, p = 3.05e-09) also exhibited the highest abundance in Orchestra samples, with Akooma and Bagoo cultivars containing greater levels compared to others. Xanthosine, another nucleoside metabolite, accumulated predominantly in Ingrid, Karacter, Manager, and Orchestra samples (η^2^ = 58.4%, p = 5.65e-11).

The relative abundances of all 74 significant metabolites influenced by cultivar effect are illustrated in a heatmap (Figure 4B). Hierarchical clustering revealed three distinct metabolite clusters. The first cluster included lipid oxidation products and lipid-like compounds, predominantly abundant in Akooma and Greenway. The second cluster included metabolites with relatively uniform intensity patterns across cultivars, including paired metabolites such as N-acylethanolamines (palmitoylethanolamide and linoleoylethanolamide), jasmonate (tuberonic acid), and phytotoxin vomifoliol. The third cluster consisted of metabolites more abundant in Kaplan (asparagine, aspartic acid, and N-benzoyl aspartic acid) and Orchestra (leucine, histidinol, methionine, and 5-methylthioadenosine).

Among the 178 SIRIUS-annotated metabolites, 105 exhibited significant differences attributed to cultivar effect. The three metabolites with the highest effect sizes were cinnamic acid conjugates with aspartic acid: caffeoyl aspartic acid (η^2^ = 96.2%, p = 7.61e-51), feruloyl aspartic acid (η = 95%, p = 2.40e-46), and coumaroyl aspartic acid (η^2^ = 93.7%, p = 2.54e-42) (Supplementary Figure 4B). These metabolites showed nearly identical patterns across cultivars, being most abundant in Orchestra and Karacter samples and absent in Ingrid samples. Isobutyryl aspartic acid (η^2^ = 85.9%, p = 7.48e-29) exhibited the highest abundance in Orchestra, Kaplan, and Ingrid samples and the lowest in Manager, Karacter, and Bagoo samples.

Soyasaponin I (η^2^ = 59.9%, p = 1.09e-11), one of the most abundant triterpenoid saponins identified in peas, showed the highest levels in Greenway, followed by Bagoo, Orchestra, Karacter, Skol, and Akooma samples. Conversely, bersimoside I (η^2^ = 51.4%, p = 7.21e-09) and pisumsaponin I (η^2^ = 42.2%, p = 2.61e-06) displayed (in total) lowest abundance in pea samples. Another saponin annotated as 12-oleanene-24-ol-3-O-Rha-Xyl-GluA-22-O-Ara (η^2^ = 59.4%, p = 1.54e-11) had higher concentrations in Akooma and Greenway cultivars. Dimethylarginine (η^2^ = 90.9%, p = 2.82e-36) was predominantly found in Akooma, Ingrid, and Greenway samples, with lower levels in other cultivars. The highest level of isovaleryl glutamic acid (η^2^ = 79.1%, p = 2.52e-22) was in Kaplan followed by Orchestra and Ingrid and absent in Bagoo and Karacter samples. Ingrid samples showed the lowest level of kaempferol 3-sophotrioside (η^2^ = 77.8%, p = 2.23e-21), while Orchestra samples had the highest abundance of this flavonoid.

Hierarchical clustering of the SIRIUS-annotated 178 metabolites identified three distinctive clusters (Supplementary Figure 4A). One cluster consisted of conjugates of aspartic and glutamic acids with cinnamic acid derivatives and short-chain fatty acids, predominantly abundant in Orchestra, Kaplan, and Karacter cultivars. Interestingly, Ingrid samples were devoid or showed significantly low levels of hydroxycinnamic acid aspartates regardless of location type. A second cluster, characterized by lipid oxidation products, exhibited the highest intensities in Akooma and Greenway samples, and included the triterpenoid saponin 12-oleanene-24-ol-3-O-Rha-Xyl-GluA-22-O-Ara. In the third major cluster, triterpenoid saponins soyasaponin I and pisumsaponin I as well as jasmonates 4,5-didehydrojasmonic acid and tuberonic acid clustered with one another. Being more abundant in Orchestra samples, a triterpenoid saponin bersimoside I and kaempferol 3-sophotrioside, another member of phenylpropanoid pathway with lowest levels in Ingrid levels, clustered with 5-methylthioadenosine and methionine.

### Tentative identification of novel indole metabolites and saponins in pea seeds using mass spectral computational approaches

Combining molecular networking-based annotation in GNPS with computational methods in SIRIUS enabled the identification of additional metabolites significantly influenced by location and cultivar effects. Specifically, a metabolite with a precursor m/z value of 469.145, annotated by SIRIUS as 1-N-β-glucopyranosyl-2-oxindole-3-acetyl-N-aspartic acid (oxIAA-Asp-N-Glc), showed significant variability across cultivars (η^2^ = 54.3%, p = 9.6e-10) (Supplementary Figure 4A-B). Although this metabolite was not identified using spectral libraries employed in our study (Metaboscape and GNPS spectral libraries), molecular networking provided meaningful chemical relationships between this metabolite and five other features (Figure 5A). OxIAA-Asp-N-Glc was linked to other features through mass differences (Δ m/z) of 162.053, 15.995, and 33.961 Da. The first two delta masses correspond to glycosylation and hydroxylation, respectively. Literature data confirms that oxIAA-Asp-N-Glc can be further glycosylated to form oxIAA-Asp-N-Glc-Glc (m/z 631.1975) and these metabolites were initially identified in tomato pericarp tissue (Östin et al., 1995). The experimental fragmentation patterns from our study matched literature data for all these features. Consecutive glucose losses from precursor ion with m/z 631.1975 yielded ions at m/z 469.1449 and 307.0923. Additional diagnostic fragment ions for OxIAA-Asp moiety were m/z 216.065 and 188.07, arising from cross-ring cleavage within the glucose ring combined with β-cleavage of the amide bond and further loss of CO, respectively. Next characteristic fragments were observed at m/z 174.054, resulting from sugar cleavage from the indole moiety combined with β-cleavage of the amide bond, and at m/z 146.059, a quinolonium fragment from the indole structure.

**Figure 5.**
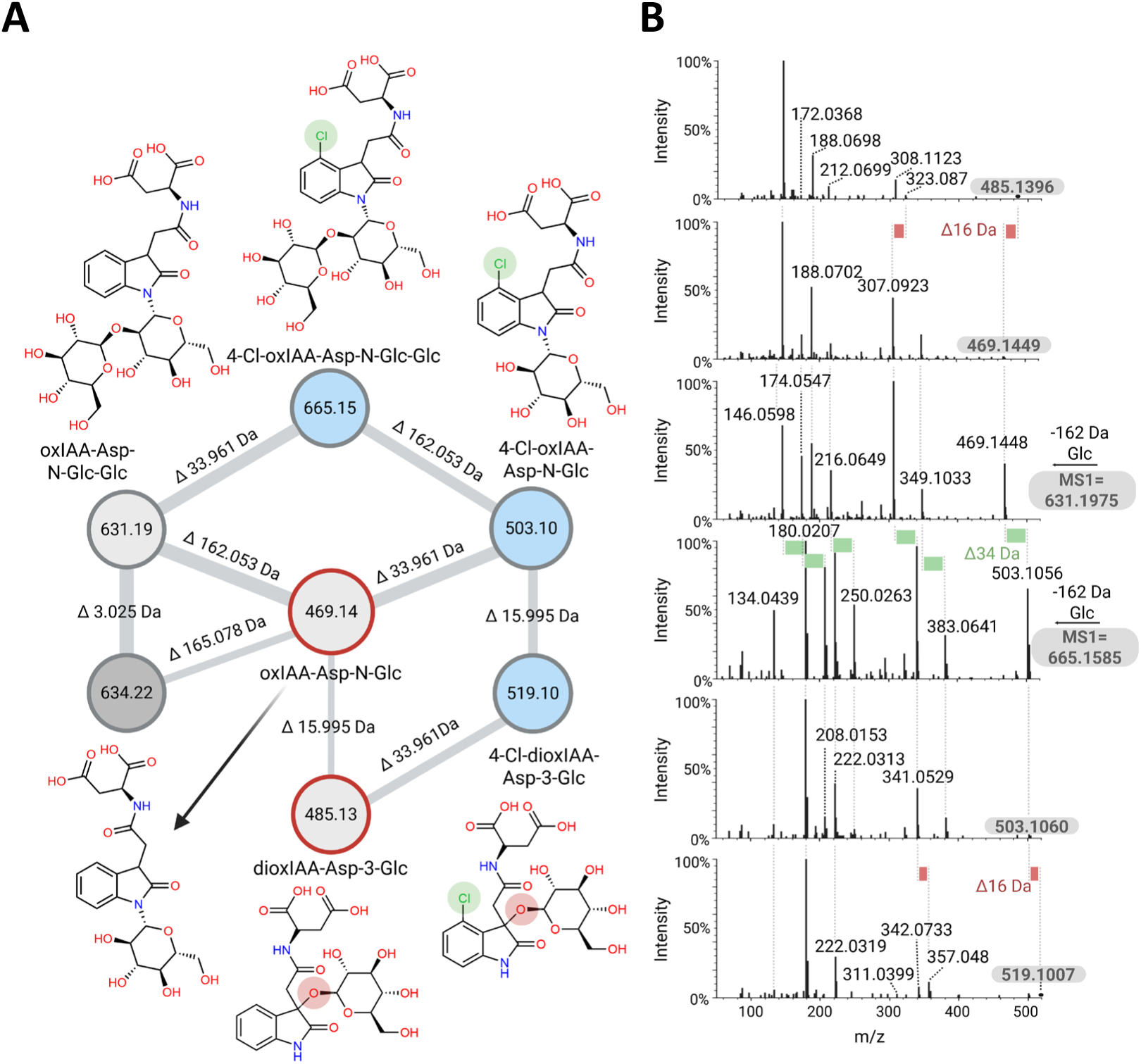
Molecular networking and SIRIUS-assisted annotation of 3-oxindole-3-acetyl aspartic acid (oxIAA-Asp) molecular family. (A) The light grey nodes indicate the MS/MS spectra of candidates were not annotated by spectral libraries, but matched with previously reported compounds in literature, the dark grey nodes – not annotated. The blue nodes mean that the putative candidates have not been annotated nor reported in literature. The nodes with red border had matches with SIRIUS annotation (B) Tandem mass spectra show diagnostic peaks of putatively annotated compounds and shifts explaining the addition of –OH and –Cl.

Regarding hydroxylation products, oxindole-3-acetic acid has been previously reported to undergo oxidation, forming 3-hydroxy-2-oxindole-3-acetic acid (Hayashi et al., 2021). This compound is subsequently glycosylated at the hydroxyl group to yield 3-O-β-glucopyranosyl-2-oxindole-3-acetyl-N-aspartic acid (dioxIAA-Asp-3-Glc, m/z 485.13), which was first identified in faba bean seedlings (Vicia faba L. cv. Chukyo), with detailed spectral characterization reported (Tsurumi & Wada, 1985). Given the absence of mass spectral evidence, specifying the precise hydroxyl group position and additional hydroxylated analogs of oxIAA-Asp-N-Glc, the metabolite at m/z 485.1396 was tentatively annotated as dioxIAA-Asp-3-Glc.

The mass difference of 33.961 Da often indicates the presence of chlorine, which we confirmed from the isotopic pattern of the precursor ion at m/z 503.1056 exhibiting characteristic isotopic peaks for the two most abundant natural chlorine isotopes, ^35^Cl and ^37^Cl. Specifically, alongside the precursor ion at m/z 503.1056 (2.3% intensity), an isotopic peak at m/z 505.1071 (1.0% intensity) was observed. Further, neutral loss of a hexose moiety (−162 Da) generated fragment ions at m/z 341.0525 (35.3%), differing by 33.961 Da from similar glucose-loss fragments from oxIAA-Asp-N-Glc, and at m/z 343.0382 (8.8%). Likewise, all other characteristic oxIAA fragment ions shifted to 33.961 Da and displayed chlorine isotope patterns, confirming chlorine substitution within the indole moiety (Figure 5B). Notably, legumes such as Lathyrus, Vicia, and Lens indeed produce 4-chloroindole-3-acetic acid, a potent auxin analogue (Engvild, 1987), though its metabolic and inactivation pathways remain largely unexplored. Therefore, the metabolite with m/z 503.1056 was tentatively annotated as 4-chloro-1-N-β-glucopyranosyl-2-oxindole-3-acetyl-N-aspartic acid (4-Cl-oxIAA-Asp-N-Glc). Molecular networking further linked this metabolite to features with mass differences of 162.053 and 15.99 Da. Examination of MS2 spectra allowed annotation of these metabolites as 4-Cl-oxIAA-Asp-N-Glc-Glc (m/z 665.1585) and 4-Cl-dioxIAA-Asp-3-Glc (m/z 519.1007). To our knowledge, these chlorinated derivatives of oxindole-3-acetic acid have not previously been reported.

4-Cl-oxIAA-Asp-N-Glc-Glc was the only chlorinated metabolite influenced by location (η = 24.5%, p = 7.22e-05), being more abundant in Hammel and Sejet than in Gamborg (Supplementary Figure 5A). All chlorinated and non-chlorinated oxIAA-Asp analogues were shown to be influenced by cultivar type in one-way ANOVA (Supplementary Figure 5B).

Similarly, metabolites annotated through SIRIUS as significant across locations and cultivars were further verified via molecular networking. Triterpenoid saponins proposed by SIRIUS lacked annotation from conventional spectral libraries, although some were tentatively dereplicated using a suspect library. For example, the metabolite with m/z 1069.558 was annotated by SIRIUS as soyasaponin βg (soyasapogenol B base + O-HexA-Hex-dHex, O-DDMP) and by the suspect library as suspect to soyasapogenol B base + O-HexA-Pen-dHex, O-DDMP (m/z 1039.547), differing by 30.011 Da (Figure 6). This mass difference corresponded to the -CH2O difference between pentose and hexose sugars, confirmed through MS2 spectral analysis showing pentose (−132 Da) and hexose (−162 Da) losses. Consequently, the metabolite at m/z 1069.558 was tentatively annotated as soyasaponin βg. This metabolite was connected to a feature at m/z 1231.601, not annotated by spectral libraries or SIRIUS, yet significantly influenced by location (η^2^ = 23.5%, p = 1.25e-04) and cultivar type (η^2^ = 46.9%, p = 3.72e-07). The 162.053 Da mass difference indicated an additional hexose unit, confirmed by MS2 spectra, revealing all characteristic soyasaponin βg ions following precursor and hexose unit losses (Figure 6). Based on available evidence, no previous studies have reported this glycosylated analog of soyasaponin βg.

**Figure 6.**
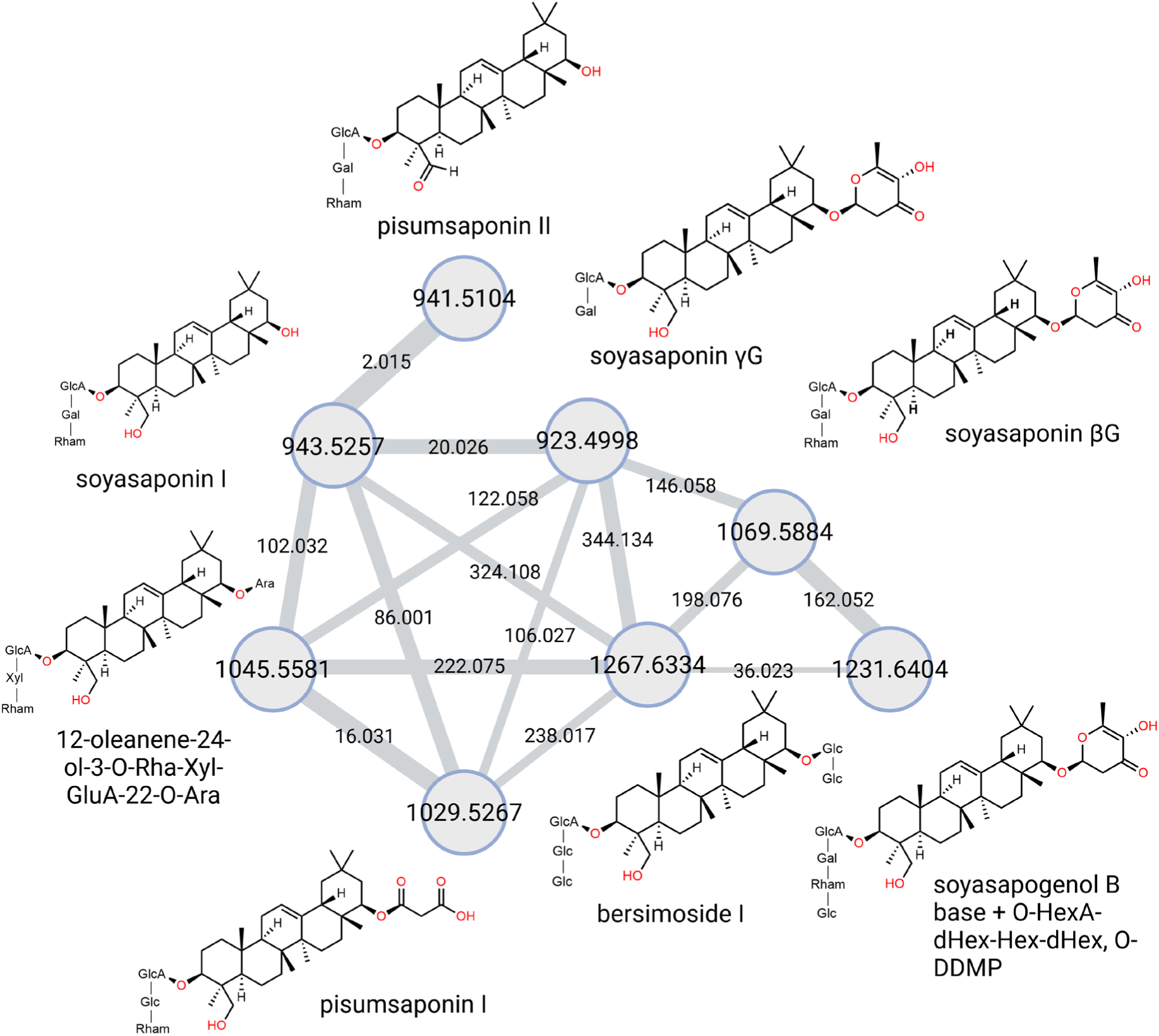
Molecular networking of pea triterpenoid saponins. Spectral libraries implemented in this study lacked reference data and were unable to annotate triterpenoid saponins. Only the combination of suspect library hits confirmed with SIRIUS annotations and manual verification of fragmentation patterns allowed for their annotations Propagation of soyasaponin βg annotation enabled the putative annotation of its analogue with mass difference 162 Da implying one more hexose unit which was confirmed in MS/MS spectra comparison.

## Discussion

Previous studies have demonstrated that peas adapt their biochemical profiles in response to both environmental conditions and genetic makeup (Mohammed et al., 2018) (Maharjan et al., 2019). Domestication and selective breeding of agronomic crops, including peas, have been reported to favor genomic characteristics enhancing growth and storage of bioactive compounds at the expense of stress resistance mechanisms. For instance, comparative analyses between wild and cultivated peas revealed that domestication typically leads to lower expression of defense-related genes and reduced accumulation of protective phenolic compounds in seed coats (Klčová et al., 2024).

Organic farming practices expose plants to unique conditions involving different nutrient availability, pest pressures, weeds, fungi and varying soil microbiomes compared to conventional farming systems. Such conditions may activate plant defense-related metabolic pathways. For instance, inoculation with nitrogen-fixing *Rhizobium* bacteria has been shown to enhance triterpenoid saponin accumulation in pea seeds, which correlates with increased resistance against fungal pathogens such as *Didymella pinodes* (Ranjbar Sistani et al., 2017). To comprehensively capture variations in complex plant metabolomes, untargeted metabolomics serves as a powerful analytical method, generating detailed chemical fingerprints useful for comparisons across varieties, treatments, and environmental conditions.

In this study, untargeted metabolomics was performed on ten pea cultivars grown at three geographically distinct sites in Denmark. The Hammel location practiced organic farming, whereas Gamborg and Sejet use conventional methods. It was found that Hammel-grown samples contained elevated levels of nitrogen-rich metabolites and non-proteinogenic amino acids, such as homocitrulline, asparagine, homoarginine, spermidine, argininosuccinic acid, and pipecolic acid, compared to those from Sejet and Gamborg. A critical distinction between organic and conventional farming systems is the source and timing of nitrogen availability. Conventional systems typically supply readily available inorganic nitrogen, whereas organic systems rely on microbial mineralization of organic matter, manure, compost, and nitrogen fixation by legumes. Additionally, organic practices promote diverse soil microbiomes, enhancing symbiotic relationships with rhizobia bacteria that convert atmospheric nitrogen (N_2_) into ammonia (NH_3_). However, excess ammonia is toxic, leading plants to rapidly assimilate ammonia into organic nitrogen-storing compounds. Generally, higher ammonium nutrition results in greater accumulation of free amino acids compared to nitrate-based nutrition (Neuberg et al., 2010). Asparagine, for instance, functions as a primary nitrogen storage and transport amino acid in peas (Neuberg et al., 2010). In pearl millet, asparagine was reported to mitigate ammonia toxicity during drought stress, serving as a nitrogen reserve that could be remobilized upon recovery of growth conditions (Kusaka et al., 2005). Additionally, Igbasan et al. demonstrated that elevated nitrogen availability significantly increases arginine content in peas (Igbasan et al., 1996). Under conditions of drought, extreme temperatures, or other abiotic stresses, plants commonly accumulate polyamines such as spermidine, which function as antioxidants and osmoprotectants (Shao et al., 2022). Pipecolic acid, a lysine metabolism product, is recognized as a signaling molecule involved in plant immunity, notably accumulating during the activation of systemic acquired resistance (SAR) in response to pathogen attack (Bernsdorff et al., 2015) (Wang et al., 2018). As organic cultivation often subjects plants to greater stress, these findings may explain the elevated accumulation of nitrogen-rich amino acids and polyamines such as spermidine in peas grown under organic conditions in Hammel.

Compared to other cultivars, Akooma and Greenway pea cultivars exhibited notably higher levels of lipid peroxidation products, likely attributable to their distinct fatty acid compositions and enzyme activities. Pea seeds primarily contain linoleic acid (C18:2) and linolenic acid (C18:3). Oxidation of these polyunsaturated fatty acids (PUFAs) by the enzyme lipoxygenase (LOX) generates aldehydes, ketones, and hydroxy-fatty acids, collectively termed oxylipins. Oxylipins are signaling molecules in plants that mediate stress responses and innate immunity. These compounds are also known contributors to the characteristic “beany” off-flavor in peas (Manouel et al., 2024). Indeed, previous research linked the levels of volatile beany compounds to both PUFA content and LOX enzyme activity. Consistent with these findings, linolenic acid was found in significantly greater abundance in Akooma and Greenway samples than in other cultivars (Figure 4A-B), possibly providing increased substrate availability for LOX-mediated oxidation. Moreover, the higher levels of 13-keto-C18:2, epoxy-C18:1, and 11-OH-C18:2 observed in Hammel samples suggest an important role of LOX-derived oxylipins in mediating plant-pathogen interactions under organic farming conditions (Blée, 2002).

The phenylpropanoid pathway, particularly the biosynthesis of hydroxycinnamic acid amides (HCAAs), was also differentially influenced by cultivar type. Higher accumulation of aspartic acid conjugates of caffeic, ferulic, and coumaric acids in Orchestra, Karacter, and Bagoo cultivars could suggest activation of this pathway in response to biotic stress and wounding (Campos et al., 2014). HCAAs can reinforce plant cell walls, providing increased resistance against microbial enzymes or directly functioning as antimicrobial agents (Liu et al., 2022). Additionally, hydroxycinnamic acids are potent radical scavengers, and their conjugation with aspartic acid might facilitate storage in less toxic forms (Bassard et al., 2010). In sensory terms, these three specific HCAAs negatively correlate with bitterness, although their unconjugated forms, such as caffeic acid, positively correlate with bitterness perception in peas (Cosson et al., 2022). Interestingly, the popular pea cultivar Ingrid completely lacked these HCAAs. A recent study has shown that wild pea genotypes have higher expression of enzymes involved in phenylpropanoid pathway than domesticated varieties, namely COMT (caffeic acid 3-O-methyltransferase) that is responsible for conversion of caffeic acid to ferulic acid (Klčová et al., 2024). Overall, transcriptomic and metabolomic results in this study demonstrated altered profile of expressed genes and metabolites, including hydroxycinnamic acids, favoring seed defense in wild (and pigmented) genotypes. Such modification may potentially be the result of targeted breeding efforts aimed at improving taste and appearance, which may explain the absence of caffeoyl, feruloyl, and coumaroyl aspartates in the widely cultivated pea variety Ingrid. However, specific physiological purpose behind the conjugation of these three hydroxycinnamic acids with aspartate has not been clearly elucidated to date. It is known that the acyl acid amido synthetases of the *Gretchen Hagen 3* (GH3) family conjugate amino acids including aspartate to phytohormones such as indole-3-acetic acid, salicylic acid, phenylacetic acid, and benzoic acids to regulate levels of their active and inactive forms (Westfall et al., 2016). But whether GH3 enzymes could conjugate hydroxycinnamic acids to aspartate for similar purposes is yet to be determined.

Furthermore, 1-N-β-glucopyranosyl-2-oxindole-3-acetyl-N-aspartic acid (oxIAA-Asp-N-Glc), annotated by SIRIUS, displayed significant cultivar-dependent variability. Expansion of this annotation through molecular networking uncovered three novel chloroauxin metabolites, which were tentatively annotated by manual interpretation of their MS/MS spectra and comparison with the fragmentation patterns of their known non-chlorinated analogues. Typically, structural elucidation of previously unknown compounds requires detailed nuclear magnetic resonance (NMR) and mass spectrometry analysis of pure, isolated, or synthesized reference compounds. Subsequent matching of spectral data and retention times from biological samples confirms the presence of candidate metabolites. Although the chloroauxins proposed here, lack direct verification through such approaches, their annotations are robustly supported by well-characterized fragmentation patterns of non-chlorinated analogues and distinct chlorine isotopic signatures. Additionally, a plant-specific mass spectral repository search (plantMASST) confirmed the presence of 4-Cl-oxIAA-Asp-N-Glc in publicly available pea metabolomics data (Supporting Figure 6), affirming its occurrence as a common pea metabolite (Gomes et al., 2024). In fact, a previous pea metabolomics study also reported two metabolites at Level 3 identification as ‘chlorinated aspartic acid derivatives’, the precursor masses of which match 4-Cl-oxIAA-Asp-N-Glc and 4-Cl-dioxIAA-Asp-3-Glc found in our study (Elessawy et al., 2021). These findings emphasize the significant utility of computational tools and molecular networking in uncovering previously uncharacterized plant metabolites as well as reusing taxonomy-informed publicly available MS data to confirm annotations.

The proposed chloroauxin metabolites are analogues of indole-3-acetic acid (IAA), a widely occurring phytohormone involved in plant growth. Recently, comprehensive insights into the biosynthesis of IAA and its reversible and irreversible inactivation pathways have been described (Fukui et al., 2022). Notably, several agriculturally significant legumes, including pea, lentil, and faba bean, produce the more potent auxin analogue 4-chloroindole-3-acetic acid (4-Cl-IAA), which plays an essential role in pod elongation (Reinecke, 1999). IAA homeostasis is maintained through reversible inactivation by conjugation with amino acids and sugars (via *GH3-IAA-Leu-Resistant1 (ILR1)* enzymes and UDP-glycosyltransferases, respectively) and irreversible catabolism via oxidation of the indole moiety (via *DIOXYGENASE FOR AUXIN OXIDATION 1 (DAO1)-ILR1)* (Hayashi et al., 2021). However, the metabolic fate of 4-Cl-IAA – whether it undergoes degradation or conjugation – remains unclear (Ludwig-Müller, 2022). Previous studies have identified 4-Cl-IAA methyl ester and its aspartate amide conjugate in pea seeds (Hattori & Marumo, 1971), and quantitative analyses have detected these compounds in young developing seeds and pericarps (Magnus et al., 1997). Moreover, a recent study demonstrated *in vitro* that chlorinated IAAs can be converted to amino acid conjugates by the GH3 auxin amino acid synthetase family, enzymes known for their role in IAA-amino acid conjugation (Walter et al., 2020). These findings suggest that 4-Cl-IAA catabolism may closely resemble the metabolic pathways of non-chlorinated IAA, involving amino acid conjugation. However, oxidative catabolism of 4-Cl-IAA has not yet been reported, and further research is necessary to elucidate the enzymes involved.

In summary, this study illustrates the impact of growth conditions, specifically conventional versus organic farming practices, and cultivar differences in the pea seed metabolome. Both multivariate and univariate analyses demonstrated that selected pea cultivars exhibited biochemical adaptations to environmental and genetic factors via alterations in amino acid, fatty acid, phenylpropanoid, and triterpenoid saponin pathways. Furthermore, variations in compounds associated with sensory off-flavors were examined. Enhancing traditional untargeted metabolomics approaches with computational annotation methods significantly expanded our understanding of location- and cultivar-specific metabolites and facilitated the identification of novel chloroauxin catabolites.

## Materials and methods

### Field trials and samples

Ten cultivars of field peas (*Pisum sativum* L.), Akooma, Astronaute, Bagoo, Greenway, Ingrid, Kaplan, Karacter, Manager, Orchestra, and Skol, were grown at 3 different geographic locations in 2022 (30 samples in total). At two locations (Sejet and Gamborg), peas were grown according to conventional principles. Sejet is a clay loam/heavy clay soil while Gamborg is a silty loam/clay loam soil. Peas were sown April 1^st^ or March 31^st^ at the Sejet and Gamborg location, respectively. Peas at the Gamborg location received 400 kg 0-4-21 synthetic NPK fertilizer per hectare at sowing. At the Sejet and Gamborg locations pesticides were applied to control weeds, fungi and pests. At the third location (Hammel, a coarse sandy clay soil) peas were grown according to the principles of organic farming (EU Regulation 2018/848). Sowing was conducted on the 12^th^ of May and no fertilizers or pesticides were applied. At maturity, ultimo August/primo September, peas were harvested and stored in aerated bags.

### Sample processing

All samples were extracted in 80% methanol (HPLC-grade, Sigma-Aldrich, Germany) in randomized order using 2 mL Eppendorf tubes. The extraction solvent contained 4 µM sulfadimethoxine (VETRANAL™, analytical standard, Sigma-Aldrich, Germany) as an internal standard to monitor LC-MS/MS system performance. Samples were vortexed (Vortex-Genie 2, Scientific Industries, USA) for 10 seconds, followed by ultrasonication (Sonorex Digitec DT 100 H, Bandelin, Germany) at 35–40 °C for 30 minutes. Samples were vortexed again for 10 seconds and centrifuged (ScanSpeed 1580R, LaboGene, Denmark) at 21,124 × g for 20 minutes at 4 °C. Subsequently, 200 µL of supernatant was transferred into glass inserts (Mikrolab Aarhus A/S, Denmark) within 2 mL glass vials (Mikrolab Aarhus A/S, Denmark).

### LC-MS/MS data acquisition

Extracted samples were randomized and analyzed using an Elute ultra-high performance liquid chromatography (UHPLC) system coupled with an impact II quadrupole time-of-flight mass spectrometer (QTOF-MS) (Bruker Daltonik, Bremen, Germany). Pooled quality control (QC) and blank samples were analyzed every eight biological samples. Chromatographic separation was achieved using a reverse-phase Intensity Solo 2 C18 column (100 Å, 2 µm, 100 mm × 2.1 mm, Bruker Daltonik) preceded by an ACQUITY UPLC BEH C18 VanGuard pre-column (130 Å, 1.7 µm, 2.1 mm × 5 mm, Waters Corporation, Milford, MA, USA), maintained at 40 °C. The mobile phase consisted of solvent A (Milli-Q water + 0.1% formic acid, Merck, Germany) and solvent B (acetonitrile + 0.1% formic acid, hypergrade LiChrosolv®, Merck, Germany) with a constant flow rate of 0.3 mL/min. The gradient elution was as follows: 0–2 min (100% A), 2–8 min (linear gradient to 100% B), 8–11 min (100% B), 11–11.01 min (return to 0% B), and 11.01–14 min (re-equilibration at 0% B).

Mass spectra were recorded from m/z 20 to 1300 at a spectral rate of 12 Hz using electrospray ionization (ESI) in positive-ion mode. The instrument was externally calibrated using a 10 mM sodium formate solution in 50% isopropanol before sample analysis and tuned with ESI-L Low Concentration Tuning Mix (Agilent Technologies, Santa Clara, CA, USA). ESI conditions were: end plate offset: 500 V, capillary voltage: 4500 V, nebulizer pressure: 2.2 bar, dry gas flow: 10 L/min, and drying temperature: 220 °C. MS conditions were: funnel 1 and 2 RF: 200 Vpp, hexapole: 50 Vpp, ion energy: 5 eV, collision energy: 5 eV, pre-pulse storage: 5 µs. Tandem mass spectra were acquired in data-dependent acquisition mode, selecting three precursor ions per scan cycle, with active exclusion after three spectra and a release period of 0.20 min. LC-MS/MS acquisition was controlled using otofControl v5.2 and Bruker Compass HyStar v5.1 (Bruker Daltonik).

### Data analysis

Data preprocessing was performed with MetaboScape 2021b (Bruker Daltonik). Feature extraction parameters included minimum features in ≥2 analyses, minimum presence in ≥3 analyses, intensity threshold of 5000 counts, and a minimum peak length of 6 spectra (4 spectra for recursive extraction). Extracted ion chromatogram (EIC) correlation was set at 0.8, considering ions: [M+H]+, [M+Na]+, [M+K]+, [M+NH4]+, [M+H-H2O]+, [2M+H]+, [M+H+H]2+, and [M]+, with [M+H-H2O]+ as a common fragment ion. Mass recalibration utilized auto-detection (Na Formate pos). Spectral annotation employed hierarchical search mode, validating annotations exclusively by MS/MS spectra with precursor m/z tolerances of 2–5 ppm, mSigma tolerances of 20–50, and MS/MS scores ranging from 900 to 600.

Feature-based molecular networking (FBMN, v2024.10.09) was conducted within the GNPS2 ecosystem (gnps2.org), applying a precursor ion tolerance of 0.05 Da, fragment ion tolerance of 0.05 Da, cosine score, and minimum matched peaks set at 0.65 and 6, respectively. Molecular networks were visualized using Cytoscape v3.10.3. Computational annotation via SIRIUS (v6.0.6) employed CSI:FingerID for molecular structure predictions, with top-ranked candidates assigned confidence scores (0 to 1) through COSMIC, which combines E-value estimation and support vector machine classification.

ANOVA-simultaneous component analysis (ASCA) (Smilde et al., 2005) was performed in MATLAB ver. R2023a (9.14.0.2337262) on auto scaled data using MATLAB scripts written by the authors. A one-way ANOVA was performed with p-values adjusted for multiple testing using the Benjamini-Hochberg procedure (FDR = 5%) in python (v.3.6) using scipy and statsmodels libraries. Hierarchical clustering heatmaps were generated using Ward’s method and Euclidean distance on row-wise max-normalized data using scipy library. matplotlib and seaborn libraries were used for visualization of the plots.

## Author contributions

S.B.E., K.H.L., and B.K., designed the research; S.A.K. and B.K. performed research; S.A.K. analyzed data; S.A.K. and B.K. wrote the paper; All authors read and approved of the final manuscript.

## Acknowledgements

We thank the AQRIFood project coordinator Christian Bugge Henriksen, University of Copenhagen, and the whole project consortium including university and industry partners. Special thanks goes to Carlsberg A/S for providing co-funding and the project partners responsible for field trials (Linda Kærgaard Nielsen, Sejet Plant Breeding and Inger Bertelsen, Innovation Centre for Organic Farming) and for sample preparation (Maria Monrad Rieckmann and Lisbeth Tina Dahl, University of Copenhagen).

## Data availability

The data supporting this article is available through www.github.com/bekzodkhakimov. Molecular networking job can be accessed at: gnps2.org/status?task=a4c1ca4f1f8e4312b9204e23948bee6f.

