## Supplementary Information for "Untargeted metabolomics reveals farming practice- and cultivar-driven modulations of pea (Pisum sativum L.) seed metabolome"

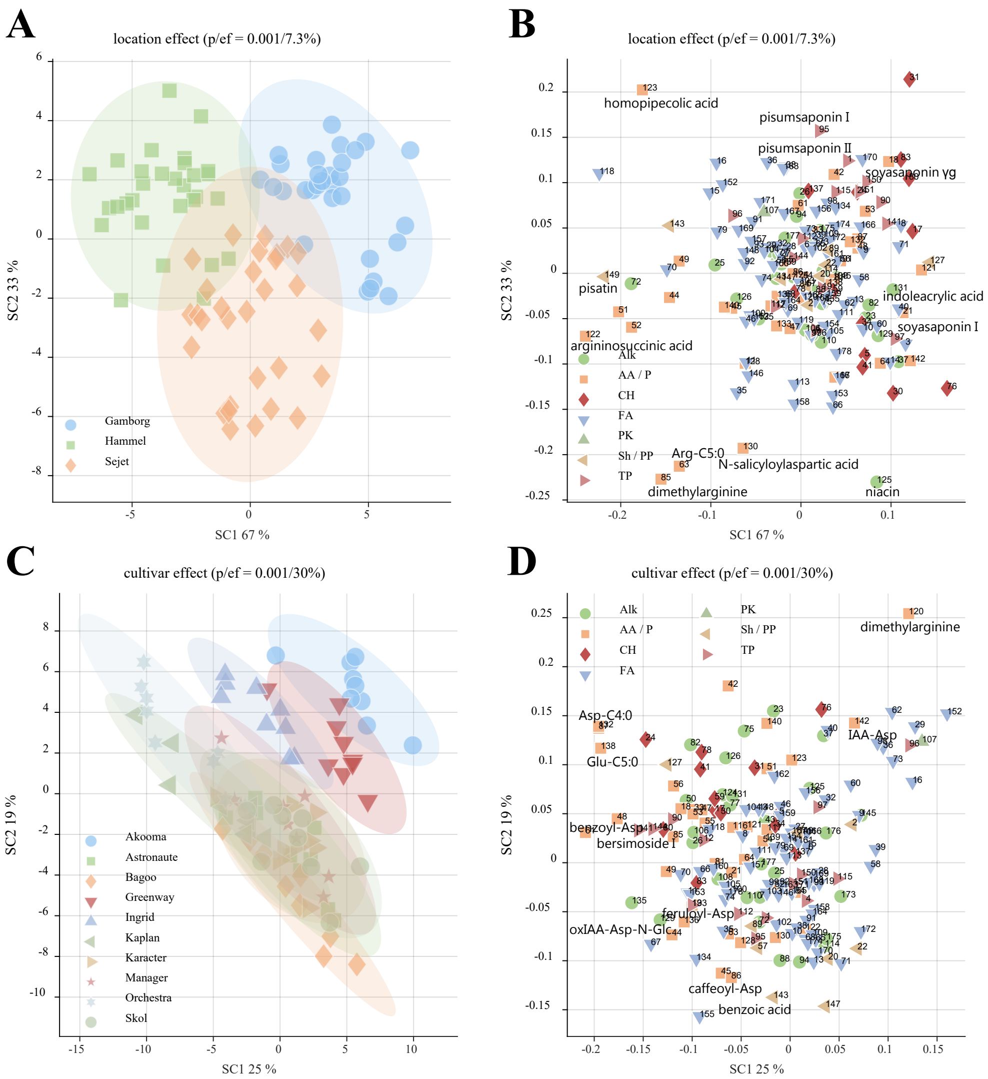


**Supplementary Figure 1.** ANOVA-simultaneous component analysis (ASCA)-based location and cultivar effect on 178 SIRIUS annotated data. **(A)** Scores plot demonstrates clearer separation of pea samples according to the three locations (variance explained = 7.3%, p-value = 0.001) (n = 3 × 30). Loadings plots show the distribution of metabolites, coloured according to biosynthetic pathways they derived from, explaining location **(B)** and cultivar **(D)** effect. **(C)** Scores plot highlights location-independent cultivar effect (variance explained = 30%, p-value = 0.001) for all pea samples (n = 10 × 9).

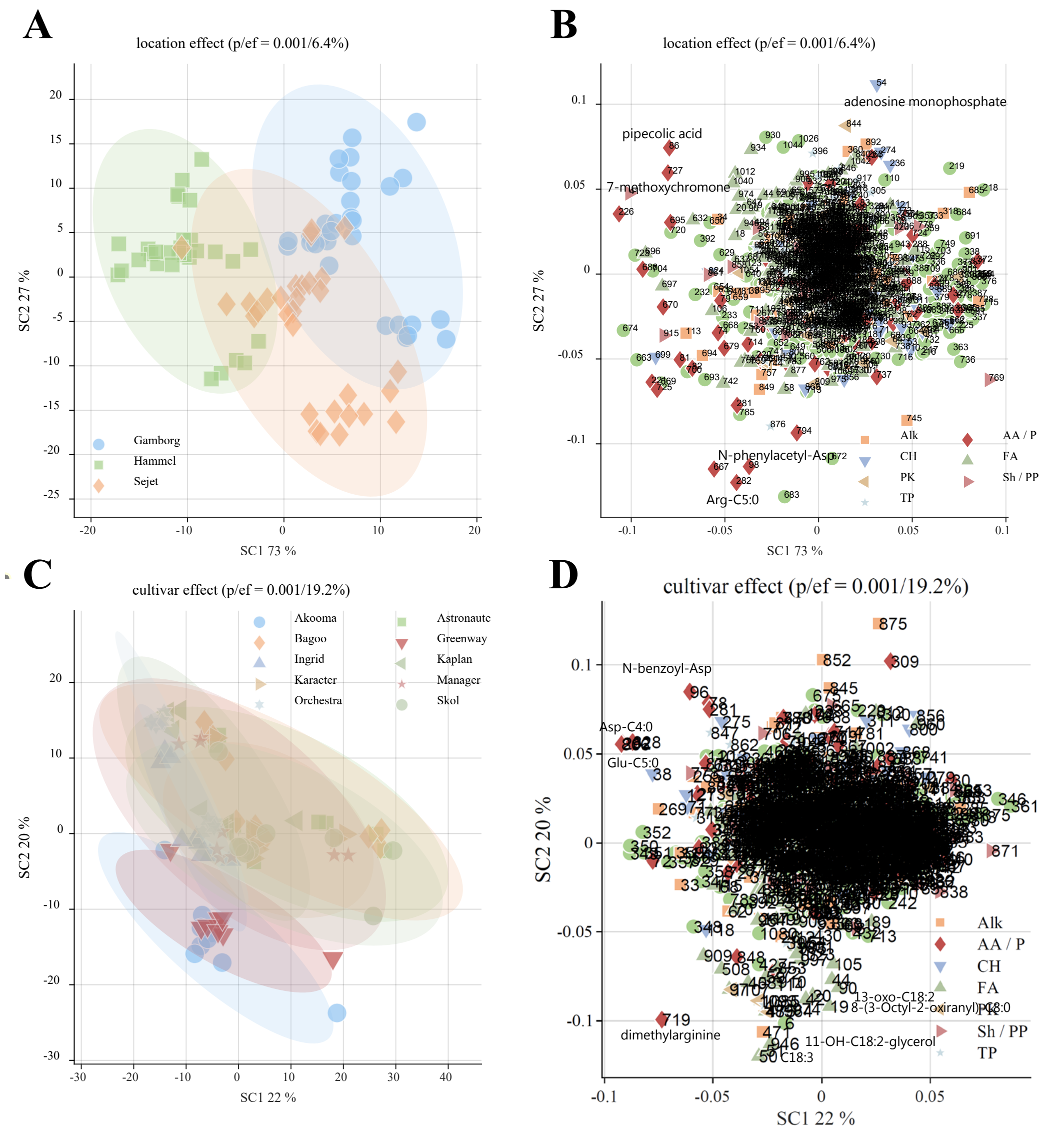


**Supplementary Figure 2.** ANOVA-simultaneous component analysis (ASCA)-based location and cultivar effect on all 1230 features. **(A)** Scores plot reiterates previous separation patterns of pea samples according to the three locations (variance explained = 6.4%, p-value = 0.001) (n = 3 × 30) except for splitting the location populations to two distinct subpopulations. Loadings plots show the distribution of metabolites, coloured according to biosynthetic pathways they derived from, explaining location **(B)** and cultivar **(D)** effect. **(C)** Scores plot highlights location-independent cultivar effect (variance explained = 19.2%, p-value = 0.001) for all pea samples (n = 10 × 9) with several cultivars also forming two separate populations.

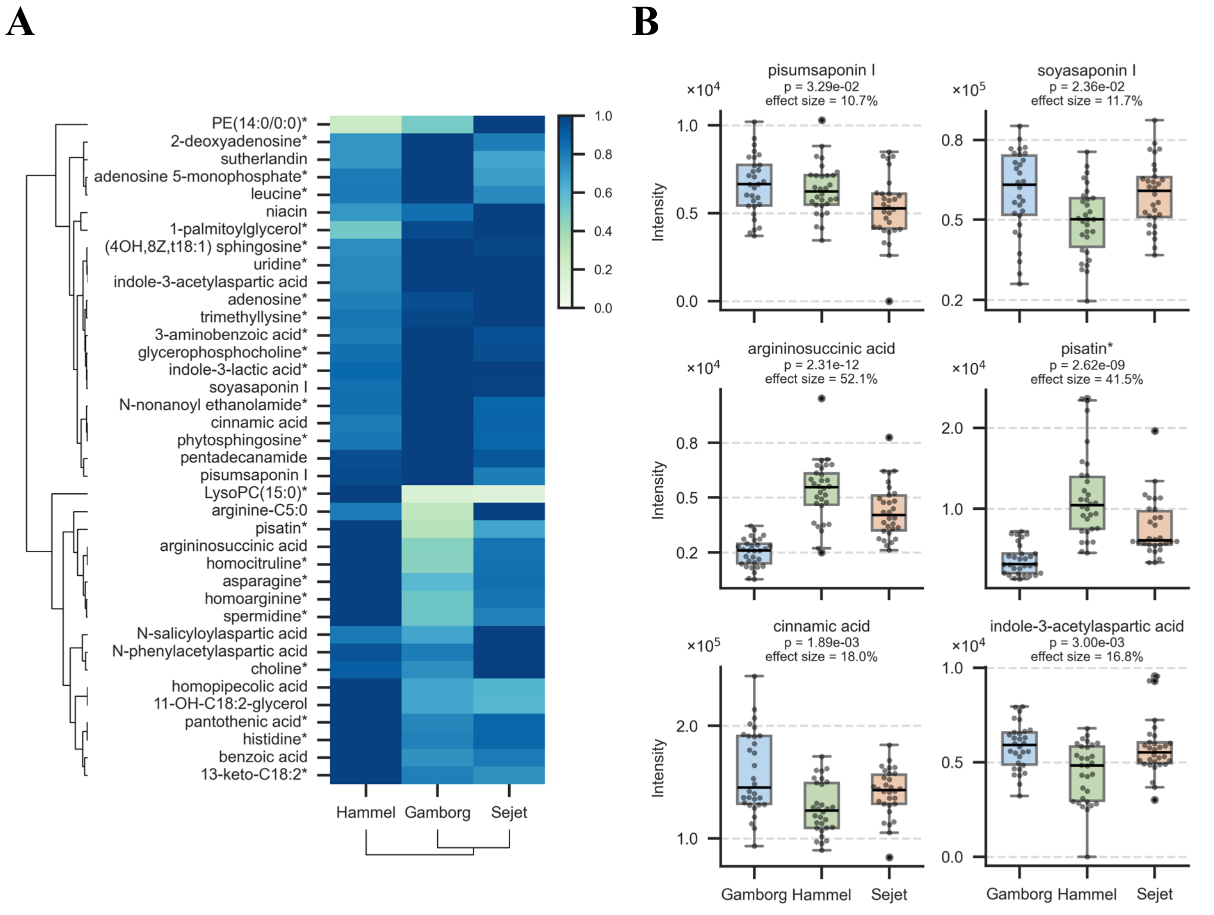


**Supplementary Figure 3.** Cultivar-independent variation of metabolites levels annotated by SIRIUS across three locations. **(A)** Hierarchically clustered heatmap of 41 metabolites identified by one-way ANOVA (FDR-adjusted p-values < 0.05 using Benjamini-Hochberg's correction for multiple testing) as significantly different between the locations. **(B)** Box plots of selected metabolites with distinct relative abundance across locations identified as significant by one-way ANOVA. * - match with spectral libraries annotations

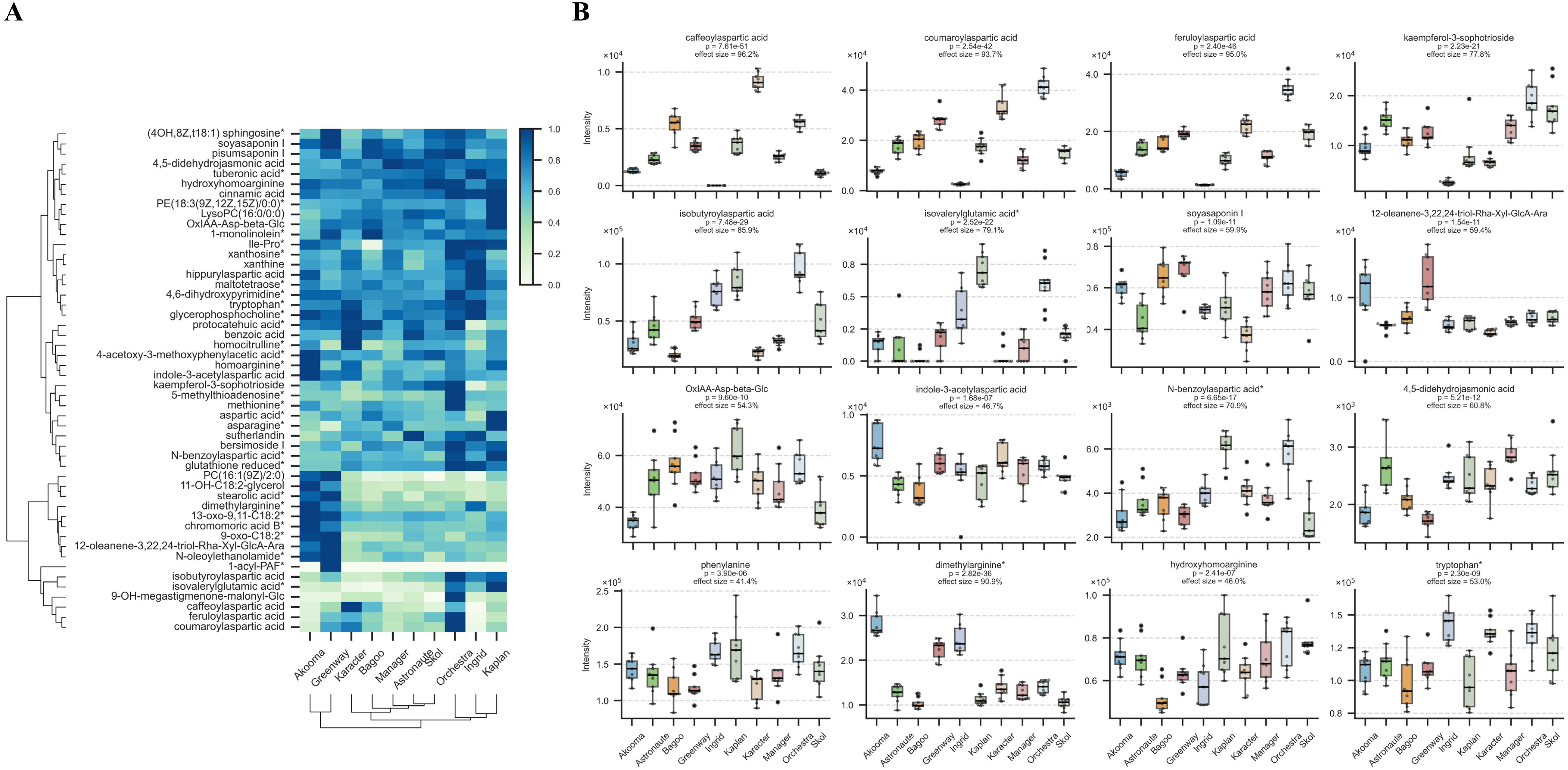


**Supplementary Figure 4.** Location-independent variation of metabolites levels annotated by SIRIUS across ten cultivars. **(A)** Hierarchically clustered heatmap of top 50 out of 105 metabolites identified by one-way ANOVA (FDR-adjusted p-values < 0.05 using Benjamini-Hochberg's correction for multiple testing) as significantly different between the cultivars. **(B)** Box plots of selected metabolites with distinct relative abundance across cultivars identified as significant by one-way ANOVA. * - match with spectral libraries annotations

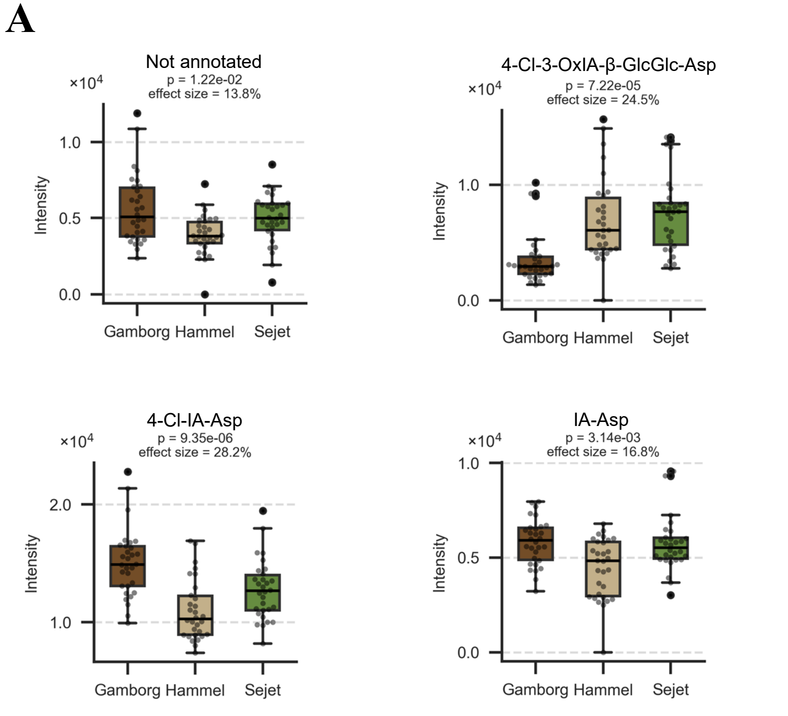


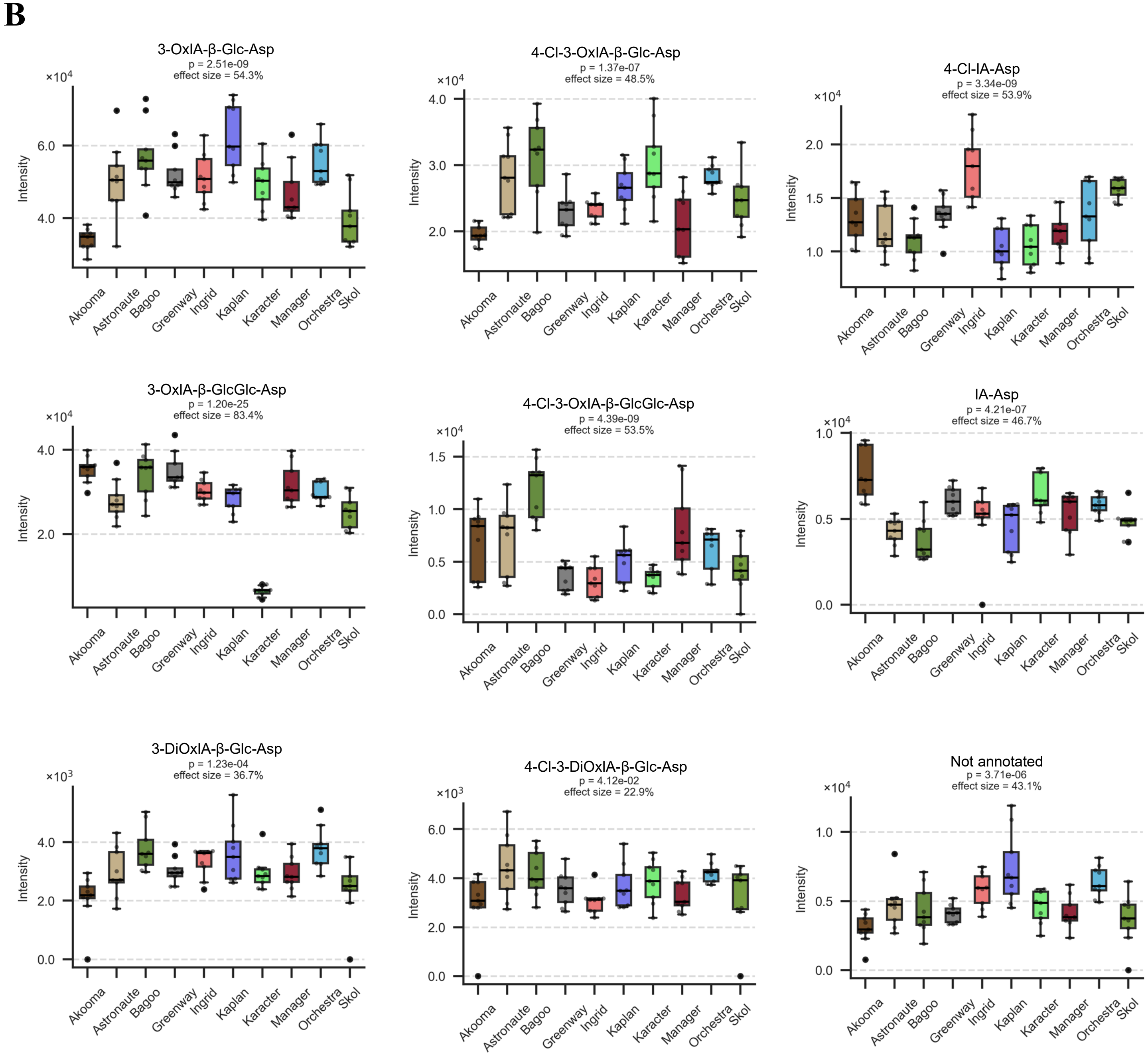


**Supplementary Figure 5.** Box plots of indole-3-acetic acid and 4-chloroindole-3-acetic acid catabolism products reflecting oxidative and conjugation with aspartic acid and glucose inactivation pathways revealed by one-way ANOVA (FDR-adjusted p-values < 0.05 using Benjamini-Hochberg's correction for multiple testing) as **(A)** location- and **(B)** cultivar specific

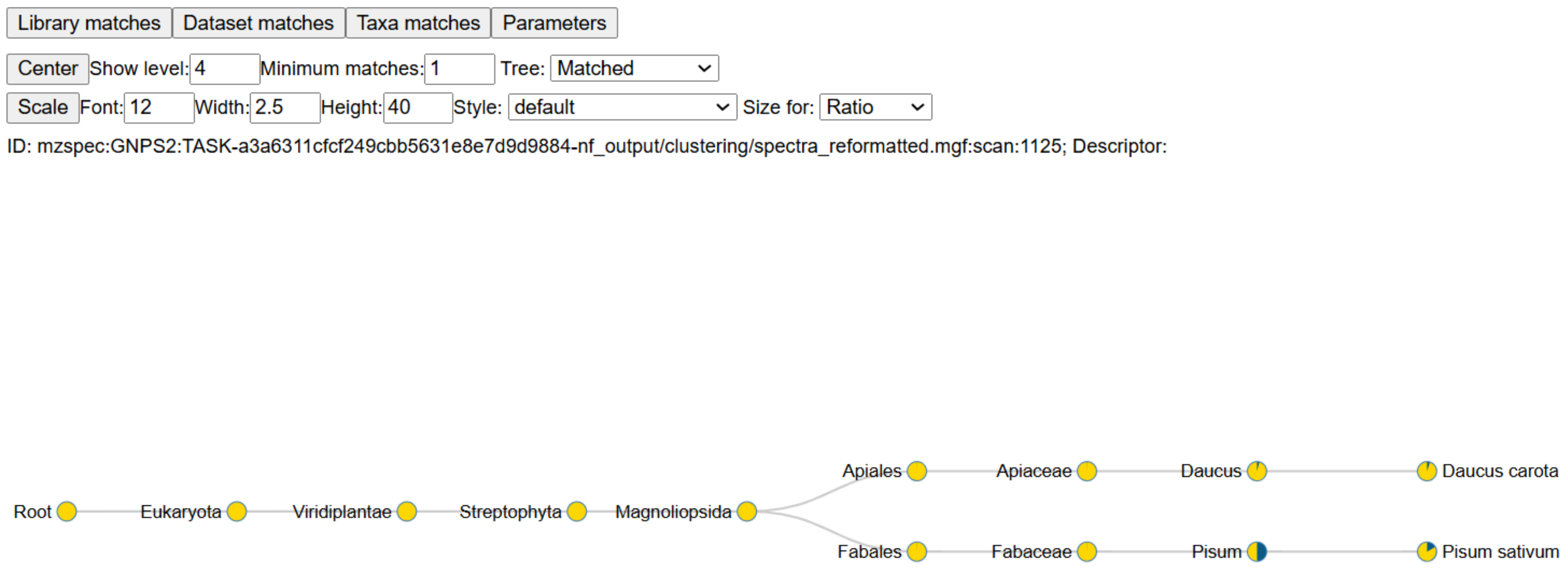


**Supplementary Figure 6.** plantMASST search of feature with m/z 503.106 tentatively annotated as 4-Cl-oxIAA-Asp-N-Glc. Precursor and fragment ion tolerance was set to 0.05 Da, cosine threshold was 0.7, minimum matched peaks was 3. In total, the accurate mass and fragmentation pattern of the queried metabolite (4-Cl-oxIAA-Asp-N-Glc) had matches in five public pea LC-MS/MS metabolomics files
